# Towards ‘Ancientbiotics’ for Biofilms: How can we bring traditional medicinal remedies out of treatise and into contemporary science?

**DOI:** 10.1101/2020.05.05.079897

**Authors:** Madhusoodhanan Vandana, Snehal Kadam, Anuradha Bandgar, Karishma S Kaushik

## Abstract

**Introduction:** Traditional medicinal remedies hold vast potential as novel antimicrobial agents, particularly for recalcitrant infection states such as biofilms. To explore their potential, it is important to bring these remedies out of ancient treatise and into present-day scientific evaluation. For traditional medical practices, this ‘development pipeline’ starts with probing treatise for potential remedies and testing them for anti-biofilm effects, or the ‘treatise to test’ phase.

**Aim:** The aim of this work is to present a primer for developing ‘ancientbiotics’ against biofilms, that focuses on the ‘treatise to test’ phase of the pipeline. Based on our approach and results, we provide insights into the considerations and challenges relevant to evaluating traditional remedies as anti-biofilm agents.

**Methodology:** We identified and reconstituted plant-based medicinal formulations from historical treatises of Indian traditional medicine, and analyzed their efficacy using widely-employed microtiter based assays, that constitute the cornerstone of biofilm studies. Measuring biomass and metabolic activity, we evaluated effects on biofilm formation and eradication of pre-formed biofilms, of *Pseudomonas aeruginosa* and *Staphylococcus aureus*.

**Results:** Based on recipes and preparation practices across several texts, and with modifications to ensure compatibility with modern scientific practices, three plant-based traditional remedies were identified and formulated in sesame oil (*Bryophyllum pinnatum, Cynodon dactylon*, and *Ocimum tenuiflorum*). We observed differential effects on biomass and metabolic activity on the biofilm formation and eradication of *P. aeruginosa* and *S. aureus*; highlighting the value of the microtiter-based assays as an initial screening tool for traditional remedies.

**Conclusion:** Through this study, we provide insights into considerations relevant to the ‘treatise to test’ phase of the ‘ancientbiotics’ pipeline, such as identifying ancient remedies, reconstituting them with present-day modifications, and using *in vitro* assay formats for evaluation. The learnings in this primer will be relevant to both contemporary scientists and practitioners of ancient medicine, and will serve as a starting point for future studies exploring anti-biofilm approaches at the interface of historical and modern medicine.

## Introduction

The first observation of biofilms can be traced to Antonie van Leeuwenhoek’s 17th century observations of ‘little animalcules’, likely to have been microbial aggregates (1). Biofilms have probably existed since the advent of microbial life, however, it was only in the late 20th century that the term was officially coined (2,3). Since then, they have gained widespread attention owing to the serious clinical and public health concerns they pose (4). Biofilms are multicellular aggregates of bacteria, attached to each other or a surface, via a self-produced extracellular matrix (5). Implicated in a range of infection states from non-healing wounds to life-threatening burn-related sepsis (6–9), biofilms are notoriously tolerant to conventional antibiotics, resulting in persistent infections and limited therapeutic options (10–13). Consequently, there is a concerted push to develop novel anti-biofilm agents, that will expand therapeutic options, and prolong the lifetime of current antibiotics.

Historical and traditional medical practices have been recognized as valuable resources of potential new antimicrobial agents (14). These practices employ natural products, often from botanical sources, thereby providing a huge diversity of active ingredients and biological targets. Further, traditional remedies are formulated using rudimentary processes that retain the crude and combined nature of the ingredients, which is possibly important for antimicrobial activity. As evident, principles that guide historical medical approaches differ from that of conventional antibiotics, which focuses on single ingredients and unique mechanisms of action (15). Therefore, prospecting ancient antimicrobial remedies as potential anti-biofilm agents (‘ancientbiotics’) could provide a paradigm shift in the treatment of biofilms.

To advance ‘ancientbiotics’ for biofilm infections, it is essential to bring traditional medicinal remedies out of esoteric medical treatise and into scientific manuscripts (16,17). This was elegantly executed in a recent series of work, where a medieval antimicrobial remedy (‘Balds’ eyesalve’ from an Anglo-Saxon text) was reconstructed using present-day ingredients, and shown to possess anti-biofilm activity against a range of pathogens (16–18). On analysis of its safety profile, the remedy showed low levels of cytotoxicity and mucosal irritation with *in vitro* and *ex vivo* assays, and no detrimental effects in a mouse wound infection model (19); building a case to explore its potential inclusion in topical therapeutics for burns and wounds. On a larger scale, this serves to highlight the vast and untapped potential of historical antimicrobial remedies as anti-biofilm agents. To fully explore this, it is imperative to enable the steady evaluation of traditional formulations for anti-biofilm potential. In the context of traditional medical practices, this ‘development pipeline’ starts with probing treatise for potential remedies and ‘screening’ them for anti-biofilm activity (‘treatise to test’ stage), following which promising agents can be taken up for in-depth evaluation.

We present a primer for developing ‘ancientbiotics’ against biofilms, focusing on the ‘treatise to test’ phase of the pipeline. For this, we identified and reconstituted plant-based formulations from historical treatises of Indian traditional medicine, and analyzed their efficacy using microtiter-based assays, that constitute the cornerstone of biofilm studies. In doing so, we have bridged knowledge and techniques separated by at least 2000 years, to pair ancient prescriptions with modern-day scientific techniques. We discuss our approach and results, and leverage them to provide insights into the considerations relevant to evaluating traditional remedies as anti-biofilm agents.

## Materials and Methods

### Mining historical texts for potential anti-biofilm agents

To identify traditional antimicrobial remedies with potential anti-biofilm activity, we referred to historical treatises of Ayurveda, an ancient Indian system of medicine. We started with the foundational texts, ‘*Charaka Samhita*’ (compendium by *Charaka*, 1st century AD (20)) and ‘*Sushruta Samhita*’ (compendium by *Sushruta*, 4th century AD (21)). Descriptions of ailments, and associated remedies, that translated (from Sanskrit) to ‘wounds’ (‘*Vrana*’), ‘wound healing’ (‘*Vranaropana*’), ‘ulcers’, and ‘non-healing ulcers’ (‘*Dushtavrana*’), were focused on, given their present-day association with biofilm infections (22–25). To identify remedies for these conditions, we referred to the foundational texts, and the ‘Ayurvedic Materia Medica’ ‘*Bhavaprakash Nighantu*’ (written by *Bhavamisra*, ∼1600 AD (26)). From these compendiums, we identified plant-based medicinal preparations recommended for ‘wounds’ and ‘non-healing wounds’. The recipes of the formulations, ingredients, proportions and preparation practices were deciphered from ‘*Sharangdhar Samhita*’ (compendium of *Sharangdhar*, ∼1300 AD (27,28)). This text focuses on the practical aspects of preparing formulations, such as base substances to be used and extraction processes.

### Preparation of traditional medicinal remedies

Based on gleanings across compendiums, the traditional formulations were prepared by ‘extracting’ ingredients from the medicinal plants into sesame oil. Remedies of each type of medicinal plant, *Bryophyllum pinnatum* (‘*Parnabeeja*’), *Ocimum tenuiflorum* (Holy basil or ‘*Tulsi*’), and *Cynodon dactylon* (Bermuda grass or ‘*Durva*’), were prepared using a mixture of plant paste, distilled water (Alliance Industries, Pune, India) and sesame oil (Prabhat teel oil, Hanuman Trading Company, India), referred to as BPE-SO, CDE-SO and OTE-SO, respectively. The medicinal plants were procured from the local market. The leaves of *Bryophyllum pinnatum* and *Ocimum tenuiflorum*, or grass of *Cynodon dactylon*, were made into a paste using a mechanised grinder. For every one part of plant paste (100 mL), 16 parts of distilled water (1600 mL) and 4 parts of sesame oil (400 mL) were used, resulting in a 1:16:4 ratio (by volume) of plant, water, and oil respectively. The plant paste, water and oil were mixed together in a wide-mouthed steel vessel to allow for water evaporation. The mixture was boiled under low heat with constant stirring till the water completely evaporated (5-6 hours), leaving the extracts from the plant paste in the sesame oil. The extract-containing oils were filtered in a single-fold cotton cloth, to remove any remaining chunks of the leaf paste, and stored at room temperature for further use. These remedies were prepared in a single batch, and all testing was performed from the same batch.

### Bacterial strains and culture conditions

The bacterial strains used in this study, *Pseudomonas aeruginosa* (PAO1-pUCP18 (29)) and *Staphylococcus aureus* (Strain AH 133 (30)) were a gift from Dr. Kendra Rambaugh and Dr. Derek Flemming from Texas Tech University, Lubbock, USA. Strains were streaked onto Luria-Bertani (LB) agar plates (Sigma) and incubated at 37°C overnight. For liquid cultures, a single colony from the plate was inoculated in LB broth (Sigma) and incubated overnight at 37°C with shaking (150 rpm).

### Effects of traditional medicinal remedies on biofilm formation

To study the effects of traditional medicinal remedies, overnight cultures of *P. aeruginosa* and *S. aureus* were diluted to a concentration of 10_7_ CFU ml_-1_ in fresh LB broth (dilutions based on optical density measured at A_595_). From this diluted culture, 10 µl was added to a clear, U-bottom non-treated 96-well plate (Corning) to give a final count of 10_5_ CFU in each well (in triplicates). Following this, 90 µl of BPE-SO, CDE-SO and OTE-SO, were added into individual wells containing bacterial culture. Controls were set up for each set of assays that included uninoculated LB broth (sterility control), and bacterial cultures in LB without the addition of the formulations (untreated control). The plates were sealed with parafilm and incubated for 24 hours at 37°C under static conditions to allow biofilm formation. After 24 hours of incubation, crystal violet (CV) and XTT assays were used to quantify the total biomass and metabolic activity of biofilms respectively (31). All assays were done with two biological replicates, each with three technical replicates, for each strain and each formulation.

### Effects of traditional medicinal remedies on biofilm eradication

To study the effects of traditional medicinal remedies on eradication of pre-formed biofilms, overnight cultures of *P. aeruginosa* and *S. aureus* were diluted to a concentration of 10_6_CFU ml_-1_ in fresh LB broth, and 100 µl of this diluted culture was added to a 96-well plate such that each well contained 10_5_ CFUs. The plate was incubated at 37°C under static conditions to allow biofilm formation on the bottom, sides and air-liquid interface of the wells. After 24 hours, wells were washed gently with 150 µl of LB broth to remove any dead or planktonic cells, taking care not to disrupt the biofilm. Following this, 100 µl of BPE-SO, CDE-SO and OTE-SO were added individually to respective wells. Controls were set up for each set of assays that included uninoculated LB broth (sterility control), and pre-formed biofilms treated with LB broth (untreated control). Plates were further incubated at 37°C under static conditions for 24 hours. To test the viability of pre-formed biofilms after exposure to sesame oil or medicinal extracts, XTT assays were performed. All assays were done with three biological replicates, each with three technical replicates, for each strain and for each formulation.

### Crystal Violet (CV) assay for total biomass quantification

The crystal violet assay is used to quantify the total biomass of the biofilm, which includes live cells, dead cells, and matrix components. Crystal violet (CV) staining was performed on biofilms as described previously (31). Briefly, a stock solution of 0.1% CV (Sigma) was made in autoclaved distilled water, filter-sterilized and stored at room temperature in an aluminium foil covered falcon tube. From the 24-hour old biofilms in the 96-well plate, supernatant media, including planktonic cells, was removed by gentle pipetting and washing thrice with 150 µl of sterile LB broth. Biofilms were then fixed with 150 µl of 100% methanol for 15 minutes in the dark at room temperature. The biomass formed was stained with 150 µl of 0.1% filter-sterilized CV solution (previously prepared) for 30 minutes in the dark at room temperature. Excess stain was removed by washing with sterile distilled water till the color of the control wells turned clear. The bound stain was solubilized using 150 µl of 100% ethanol and mixed properly. From this, 100 µl was transferred to a new well and absorbance was measured with a plate reader (SpectraMax M5 ELISA plate reader) at A_595_.

### XTT assay for biofilm metabolic activity

The XTT assay is used to measure the metabolic activity of the biofilm (viable cells), and is based on the principle that in the presence of viable cells, the tetrazolium salt XTT is broken down into a soluble colored compound (formazan). As a result, the color formed is directly proportional to the viable cell mass in the biofilm. To measure the metabolic activity of the biofilms, the XTT assay was performed as described previously (31). A stock of XTT solution (Invitrogen) of 1 mg mL_-1_ was prepared in autoclaved distilled water, filter-sterilized and stored at 4°C till use. A stock of Menadione (SRL Chemicals) of 7 mg ml_-1_ was prepared in acetone (SRL Chemicals), filter-sterilized and stored at 4°C till use. Supernatant media, including planktonic cells, was removed by gentle pipetting and washing once with 150 µl of LB broth. Menadione was further freshly diluted in a 1:100 ratio in autoclaved distilled water, and a fresh solution of LB:XTT:Menadione in 79:20:1 ratio was prepared for each experiment. This solution (150 µl) was added to each well and incubated for 4 hours at 37°C in the dark under static conditions. After incubation, 100 µl from each well was transferred to a new well and absorbance read at A_492_.

### Statistical analysis

All assays were done with at least two biological replicates (each with three technical replicates) for each strain and for the medicinal formulations. The data was plotted as box and whisker plots. All statistical analysis was performed using GraphPad Prism 8. Data was checked for normality using the Shapiro-Wilk normality test. A one-way ANOVA with Dunnett’s multiple comparison was performed to test for significance. A p-value of <0.05 was considered significant. The effects of the remedies were expressed as a percentage reduction compared to the untreated and reported as a range.

## Results

### Historical Indian medical treatises provide descriptions of traditional antimicrobial remedies with potential anti-biofilm activity

With origins in the Indian subcontinent, Ayurveda is a traditional system of medicine, with historical compendiums of practices and preparations built over 2000 years (Fig. 1). Across the foundational treatises, ‘*Charaka Samhita*’ (compendium by *Charaka* (20)) and ‘*Sushruta Samhita*’ (compendium by *Sushruta* (21)) the management of wounds (‘*Vrana*’) and chronic, non-healing wounds (‘*Dushtavrana*’), have received substantial attention, with evidence of treatment with plant-based remedies (32). The compendium of *Sushruta* has several chapters and remedies dedicated to the management of wounds, which is not surprising given that it is a treatise on ancient surgery (33,34) (Fig. 2A, Supp. Table 1). In the compendium, wounds are described to be of two types, ‘body’ wounds (*‘Sharir Vran’*), referring to wounds as a consequence of disease states, and external wounds (*‘Agantu Vran’*) resulting from trauma, burns, and animal bites (Fig. 2A, Supp. Table 1). Today, it is well-established that bacterial infections, in the form of biofilms, constitute the single-most important cause of non-healing wounds (23), and we found evidence of non-healing wounds in the compendium of *Sushruta*, translated as the ‘germination of maggots/worms due to flies on the wound can cause pain, swelling and bleeding’ (Fig. 2B, Supp. Table 1). It is very likely that in this chronic state of tissue damage or necrosis, bacterial infection of the wound will occur and prevail (35). The compendium also describes the principles of management of non-healing wounds (‘*Dushtavrana*’) with the practice of washing the wound with a decoction of oil cooked with medicinal herbs, including ‘*Surasa*’ or *‘Tulsi*’ (Holy Basil) (Fig. 2C, Supp. Table 1). In the Ayurvedic Materia Medica (‘*Bhavaprakash Nighantu*’ (26)), the medical benefits of sesame oil are elucidated in detail, including its recommended use in medicinal oil formulations (Fig. 2D, Supp. Table 1); sesame oil is widely-used by Ayurvedic practitioners even today. Across several compendiums, we identified three medicinal plant-based formulations recommended for wound healing (‘*Vranaropana’), Bryophyllum pinnatum* (‘*Parnabeeja*’), *Cynodon dactylon* (Bermuda grass or ‘*Durva*’) and *Ocimum tenuiflorum* (Holy basil or ‘*Tulsi*’) (Fig. 2E,F,G, Supp. Table 1) All three medicinal plants have well known antimicrobial properties (36–38). The preparation practices to reconstitute these plant ingredients in sesame oil were determined from ‘*Sharangdhar Samhita*’ (Fig. 2H), which recommends that to prepare decoctions of plant-based medicinal oil, the oil component should be 4 times that of the plant paste, and the ‘liquid’ component (referring to water or milk) should be 4 times that of the oil component, resulting in a ratio of 1:16:4 for plant paste, water and oil respectively. It is unclear if this refers to measurement by weight or volume, and we decided to use volume. The recipe also calls for boiling the components together to extract the plant ingredients into the sesame oil.

**Figure 1:**
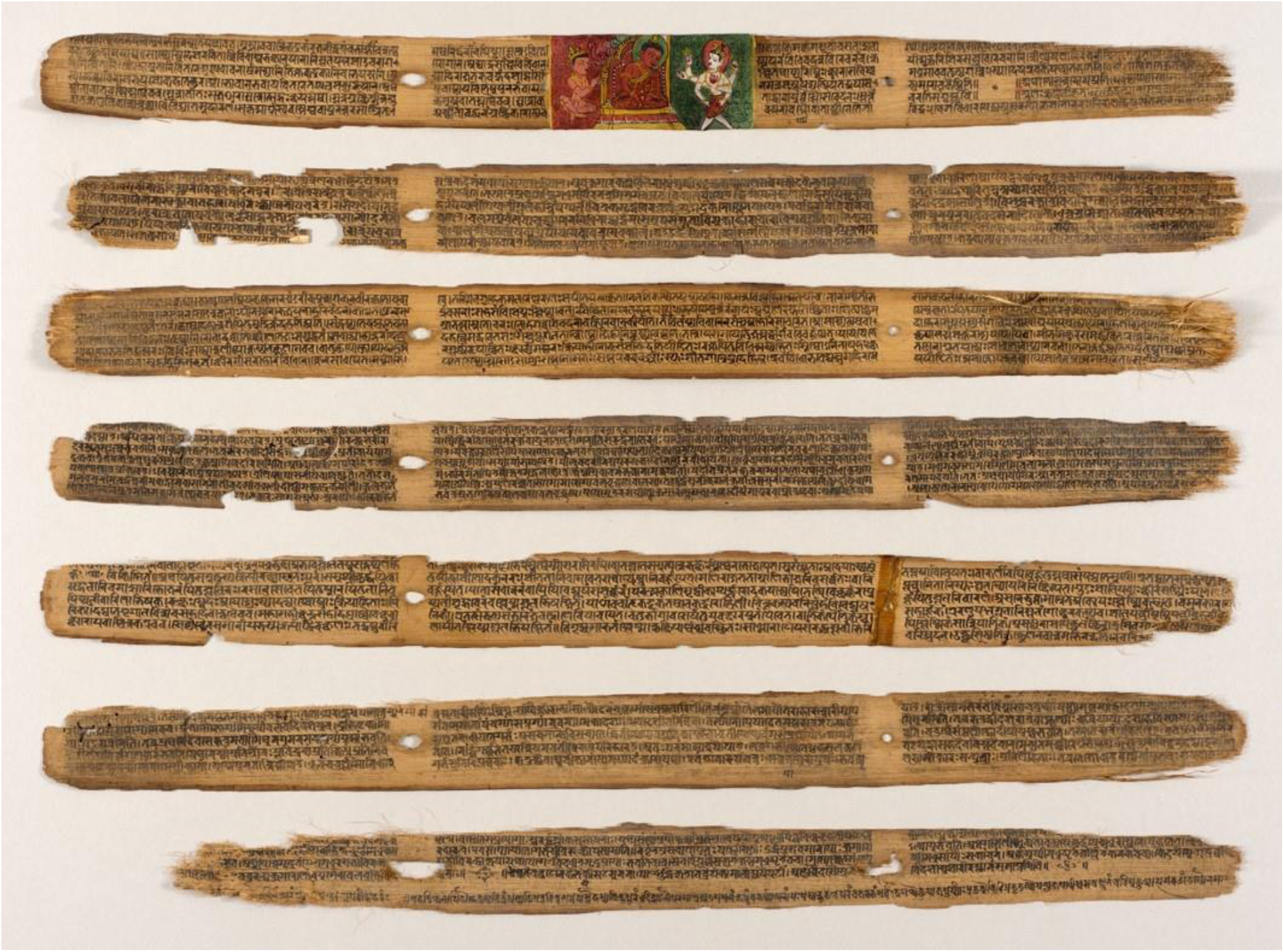
*Sushruta-Samhita*, a fundamental text of traditional Indian medicine (Ayurveda) written on palm leaves. Found in present-day Nepal, the text is dated 12_th_-13_th_ century, while the art is dated 18_th_-19_th_ century. Displayed at the Los Angeles County Museum of Art (LACMA). Reproduced in accordance with guidelines (https://www.lacma.org/).

**Figure 2:**
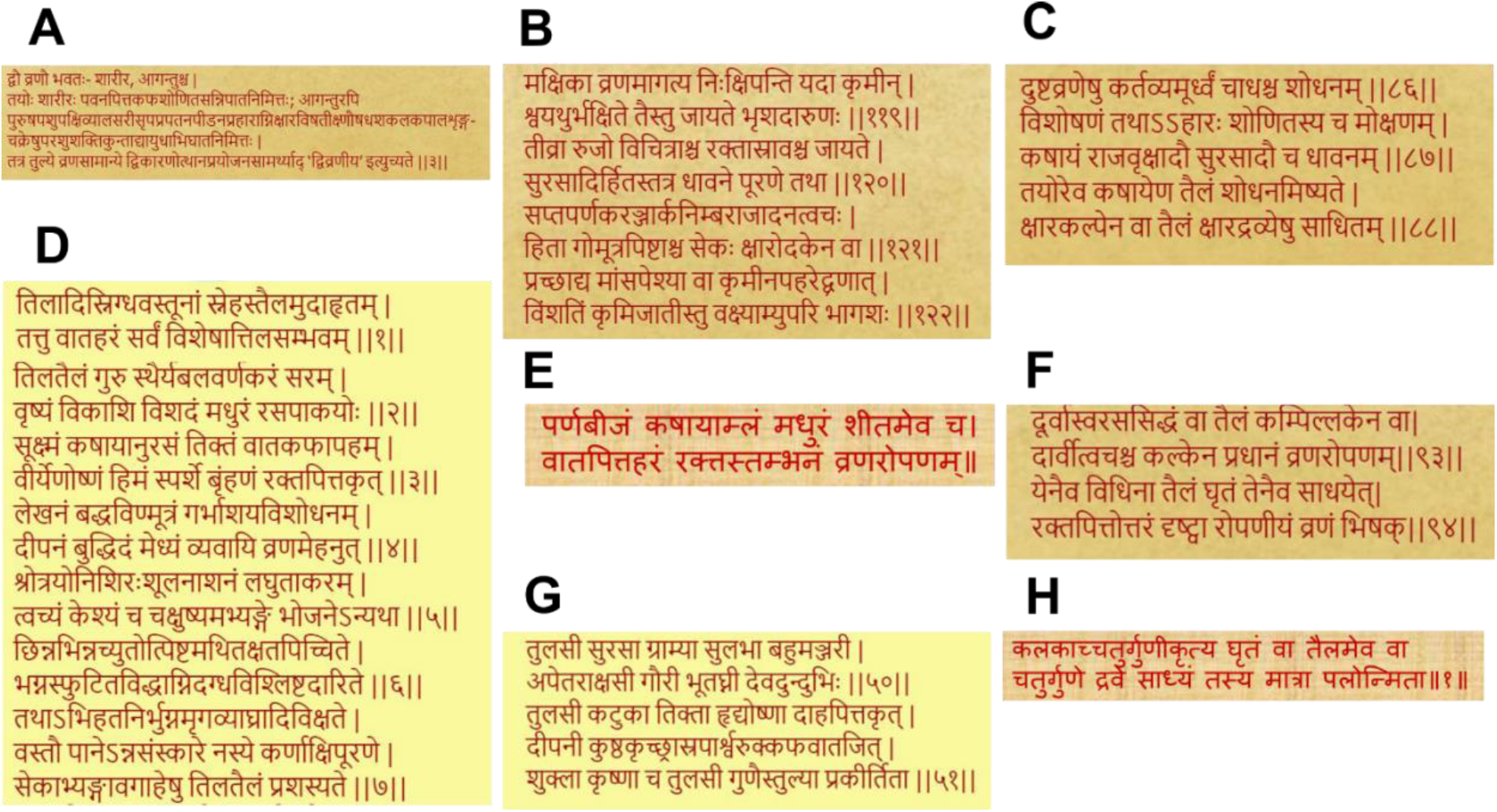
Gleanings across historical medical treatises of Ayurveda (A) Description of types of wounds in *Sushruta Samhita* (B) The problem of infected wounds described in *Sushruta Samhita* as ‘germination of maggots/worms due to flies sitting on the wound’ (C) Principles of treatment of wounds and infected wounds in *Sushruta Samhita* that call for the use of medicinal oils (D) Recommendation of the use of sesame oil for medicinal formulations in the Indian Materia Medica or *Bhavprakash Nighantu* (E) Use of *Bryophyllum pinnatum* or ‘*Parnabeej*’ for wound healing in ‘*hravyagun Vignyan* (Ayurvedic text) (F) *Cynodon dactylon* or ‘*Durva*’ for wound healing in *Charak Samhita* (G) Medicinal roles of *Ocimum tenuiflorum* or ‘*Tulsi*’ in *Bhavprakash Nighantu* (H) Preparation practices of herbal medicinal oils with four-fold ratio of components, plant paste, oil and water, from *Sharangdhar Samhita*.

### Choice of microtiter biofilm assays to test traditional antimicrobial remedies

The anti-biofilm potential of reconstituted traditional remedies was tested under *in vitro* conditions with microtiter-plate based biofilm assays (39). A mainstay of biofilm studies, these assays are widely-used, rapid, accessible, affordable, relatively easy to perform, and use commonly available materials. Further, a set of minimum information guidelines have been recently published for these assays (40). This is intended to ensure reproducibility and comparability, and we have followed these guidelines for testing and analyzing results. We evaluated the effects of the remedies on both biofilm formation and eradication of pre-formed biofilms. We measured total biomass using the crystal violet assay, and metabolic activity of the biofilms using a colorimetric assay with the tetrazolium dye ‘XTT’.

### Choice of microbial pathogens to test traditional antimicrobial remedies

Given the established association of non-healing, chronic wounds and biofilm infections (41,42), we identified and reconstituted traditional antimicrobial remedies with historical applications in the context of wounds. Today, chronic wound biofilm infections are most commonly caused by bacterial pathogens *Pseudomonas aeruginosa* and *Staphylococcus aureus* (43). We, therefore, evaluated the anti-biofilm effects of the traditional remedies on these two pathogens.

### Effects of traditional medicinal remedies on biofilm formation

To evaluate effects on the formation of biofilms, reconstituted formulations were added to bacterial cultures of *P. aeruginosa* or *S. aureus*, which were then allowed to grow under static conditions. After 24 hours, we measured total biomass of biofilms (with the crystal violet assay) and the metabolic activity of the biofilms, to indicate viable cells (with the XTT assay). As shown in Fig. 3A, in the presence of BPE-SO, CDE-SO and OTE-SO, *P. aeruginosa* showed significantly decreased biofilm formation (seen as reduced total biomass with the crystal violet assay) (p<0.001 for BPE-SO, p<0.0001 for CDE-SO and OTE-SO), when compared with untreated biofilms (UT). The reduction in the total biomass formed in the presence of the three traditional remedies was quantified to be 74-80%, 74-88% and 72-85% for BPE-SO, CDE-SO and OTE-SO respectively. This corroborates with results in Fig. 3C, that shows a reduction in biofilm metabolic activity, indicating less viable cells (XTT assay), in the presence of BPE-SO, CDE-SO and OTE-SO (75-86%, 71-79% and 49-79% reduction respectively). On the other hand, as seen in Fig. 3B, none of the three remedies, BPE-SO, CDE-SO and OTE-SO showed a significant effect on the biomass formation of *S. aureus* biofilms. However, in Fig. 3D, when treated with BPE-SO and CDE-SO there was a significant reduction in the metabolic activity of *S. aureus* biofilms (p<0.0001 and p<0.001 respectively), an effect not observed with OTE-SO. The BPE-SO and CDE-SO biofilms were quantified to have 39-60% and 15-32% reduction of metabolic activity respectively.

**Figure 3:**
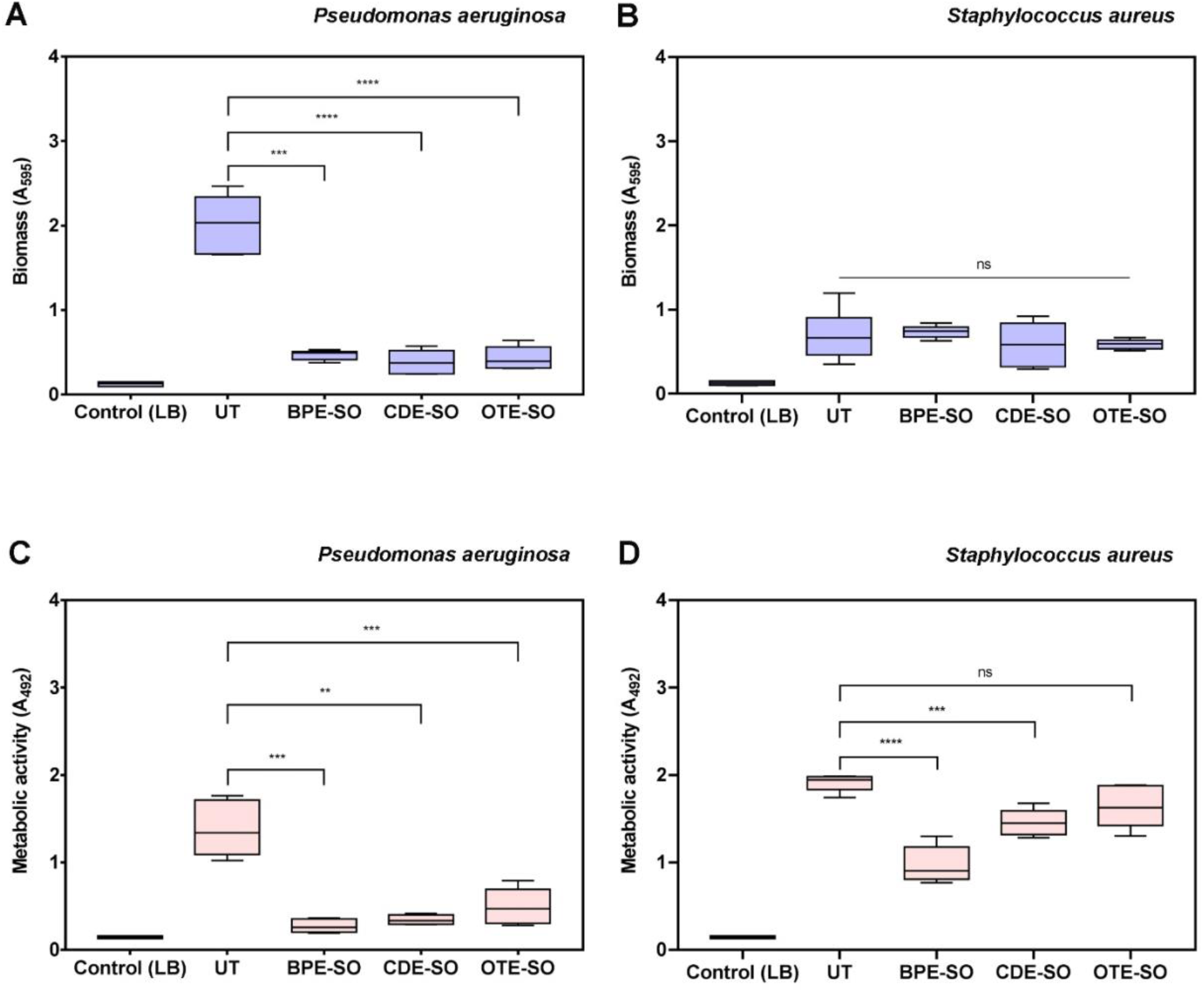
Effects of traditional medical remedies on the formation of biofilms. (A and B) Total biomass quantification (crystal violet assay) and (C and D) Quantification of metabolic activity (XTT assay) of *P. aeruginosa* and *S. aureus* biofilms formed in the presence of traditional medicinal formulations (LB-Luria-Bertani medium, UT-untreated biofilm, BPE-SO-*Bryophylum pinnatum*; CDE-SO-*Cynodon dactylon*, OTE-SO-*Ocimum tenuiflorum*; all in sesame oil(SO)). Significance was tested using a one-way ANOVA with Dunnett’s multiple comparison for the parametric data. p value ≤ 0.05 is considered statistically significant; * p< 0.05; ** p< 0.01; *** p< 0.001; and **** p<0.0001; versus untreated control, and formulations). Two biological replicates (each with three technical replicates) were performed for all assays.

### Effects of traditional medicinal remedies on eradication of pre-formed biofilms

To evaluate effects on eradication of pre-formed biofilms, reconstituted traditional remedies were added to 24-hour old biofilms of *P. aeruginosa* or *S. aureus*. After 24 hours, the metabolic activity of the biofilms was measured with the XTT assay; a critical read-out for efficacy against pre-formed biofilms is the ability to reduce the viable cell count. As shown in Fig. 4A, treatment with BPE-SO and CDE-SO were seen to reduce the viable cell mass of pre-formed 24-hour *P. aeruginosa* biofilms (p<0.01), quantified as 52-68% and 45-64% reduction respectively.

**Figure 4:**
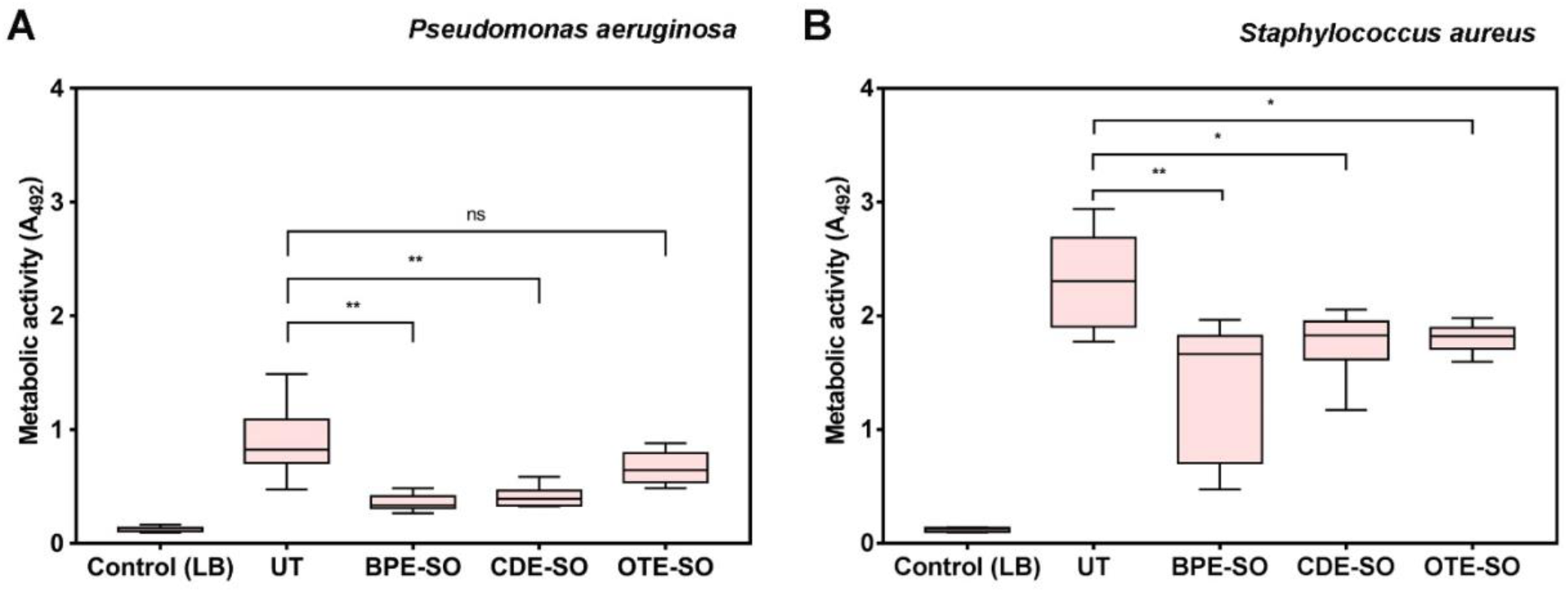
Effects of traditional medical remedies on eradication of pre-formed biofilms (24 hours old). (A and B) Quantification of metabolic activity (XTT assay) of *P. aeruginosa* and *S. aureus* pre-formed biofilms treated with traditional medicinal formulations (LB-Luria-Bertani medium, UT-untreated biofilm, BPE-SO-*Bryophylum pinnatum*; CDE-SO-*Cynodon dactylon*, OTE-SO-*Ocimum tenuiflorum*; all in sesame oil (SO)). Significance was tested using a one-way ANOVA with Dunnett’s multiple comparison for the parametric data. p value ≤ 0.05 is considered statistically significant; * p< 0.05; ** p< 0.01; *** p< 0.001; and **** p<0.0001; versus untreated control, and formulations). Three biological replicates (each with three technical replicates) were performed for all assays.

When pre-formed *S. aureus* biofilms were treated with BPE-SO, CDE-SO and OTE-SO for 24 hours, a significant reduction in the viability of biofilms was seen with BPE-SO (19-68%, p<0.01), and a lesser effect was observed with CDE-SO and OTE-SO (13-35% and 17-27% reduction respectively, p<0.05).

### Analysis of results based on minimum information guidelines for biofilm assays

Based on previously published guidelines (40), we have stated our experimental designs and protocols providing details of strains, reagent volumes, staining protocols, and types of microplates. According to the guidelines, the data was tested for normality. A description of the tests performed to obtain significance values, and the software used for this analysis are mentioned. As recommended, box and whisker plots have been used to represent data.

## Discussion

We have spanned medical and technical knowledge from the pre-2nd century CE to the present-day, to evaluate the anti-biofilm effects of traditional formulations, buried deep in historical medical treatises, using contemporary scientific assays and analysis. Based on our approach and results, we shed light on the considerations and challenges relevant to evaluating traditional medicinal remedies as anti-biofilm agents, focusing on the ‘treatise to test’ phase of the development pipeline (Fig. 5).

**Figure 5:**
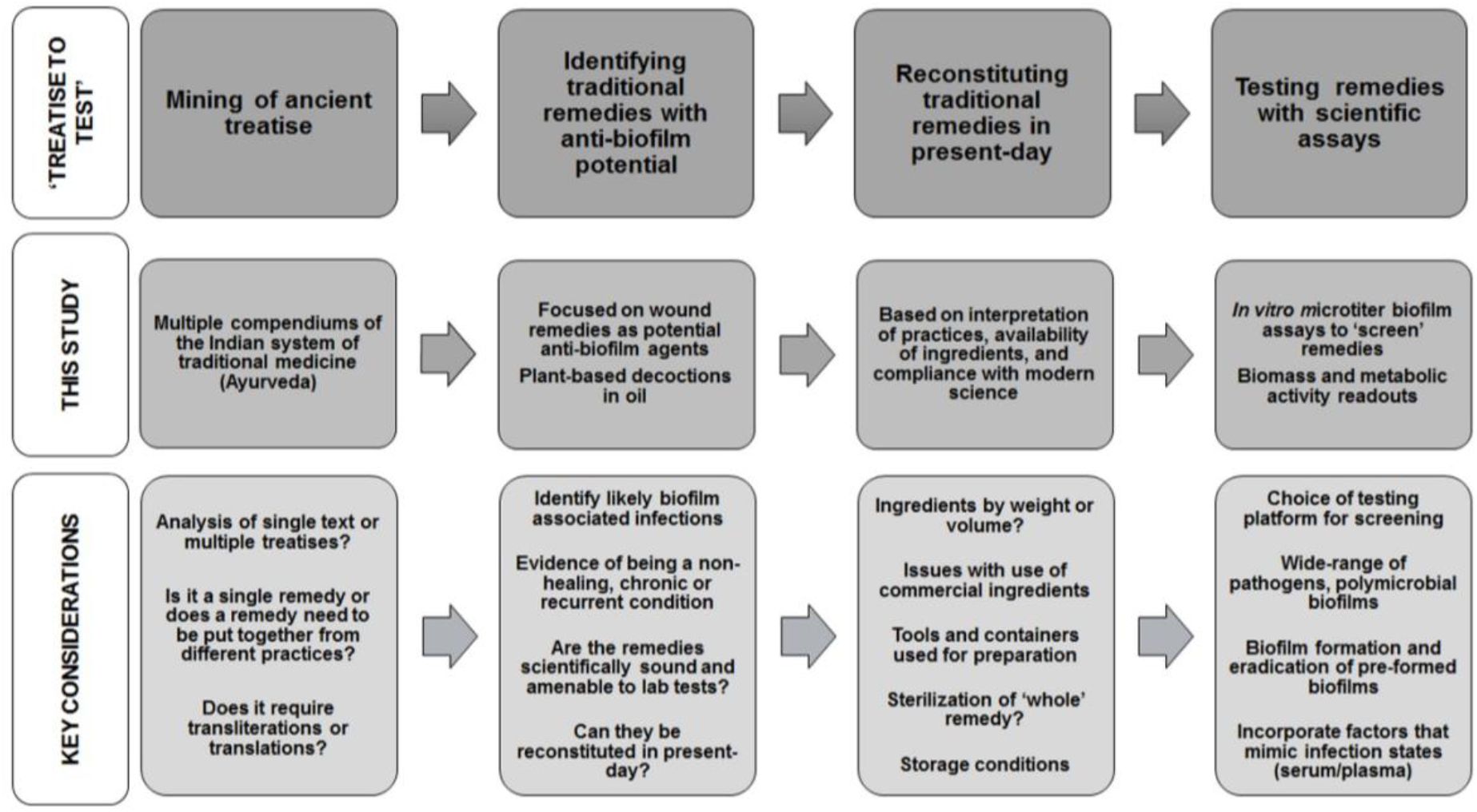
Common themes and key considerations in the ‘treatise to test’ phase of developing ‘ancientbiotics’ as anti-biofilm approaches.

### Identifying traditional remedies with anti-biofilm potential from historical treatises

Ancient texts will not explicitly include the term ‘biofilm’ (or a translated term), therefore the context of medicinal applications in the treatise was used for insights into the infection state. Given that microbial biofilms are widely implicated in non-healing wounds (23), we focused on identifying traditional wound remedies that could be explored as potential anti-biofilm agents. On similar lines, other biofilm infection states, such as eye (blepharitis or stye (44)) or ear infections (chronic suppurative otitis media (45)), could also be explored.

### Evidence of remedies being scientifically sound and testable in the laboratory

Based on findings across compendiums, we reconstituted three traditional plant-based remedies in sesame oil, *Bryophyllum pinnatum* or *‘Parnabeeja’, Ocimum tenuiflorum* or *‘Tulsi’, Cynodon dactylon* or *‘Durva’*. Sesame seed oil (*Sesamum indicum*) is known to possess antimicrobial properties (46–48), and is widely used as an ‘oil pulling’ agent to combat dental plaque formation (49–52). In an *in vitro* model of oral infection with saliva-coated microtiter plates, sesame oil displayed antibacterial activity against *S. mutans* biofilms. Further, organic oils have been shown to disrupt the formation of mature biofilms, causing cell detachment and damaged cell morphology. These effects could explain the use of sesame oil in ancient medicinal formulations. *Bryophyllum pinnatum* (53–56), *Cynodon dactylon* (57–59), and *Ocimum tenuiflorum* (36,60), possess well-known medicinal properties, and are used for a variety of ailments in traditional medicine. In ancient Indian medical treatise, we found strong evidence of their use in wound management, including chronic wounds (Fig. 2E, F, G, Supp. Table 1). Leaf extracts of these plants (methanol or ethanol-based) have been reported to have *in vitro* antimicrobial efficacy against *P. aeruginosa* and *S. aureus*, using disc-diffusion assays (54,58,60–62). Therefore, these three plant-based oil formulations were determined to be scientifically sound and testable. This is important to consider as other proposed treatments for infected wounds in treatises, such as dietary modifications, emetics and purgatives, cannot be tested in the laboratory and are out of the purview of our work.

### Reconstituting traditional remedies in present-day

To reconstitute the plant-based remedies in sesame oil, we followed preparation practices as faithfully as possible; however, certain modifications were necessary based on the availability of materials, and to ensure compliance with modern scientific practices. Based on a historical reference, the plant paste, water and oil were recommended to be used in a ratio of 1:16:4, however, it is unclear if this refers to weight or volume. The unit of measurement in traditional Indian medicine is described as ‘weight of a seed’ or ‘enough to fill your hand or palm’, the latter possibly indicating volume. We used volume to measure all components. This is an important initial consideration, and one way to overcome the lack of clarity could be to prepare different versions of the remedy.

### Use of commercial ingredients

The recipes call for ‘*teel oil’*, or sesame seed oil, which in ancient times is likely to have ‘been made from scratch’, just as the plant pastes. We did not have access to the equipment or familiarity with the processes needed to ‘press’ oil from sesame seeds, and hence used sesame oil procured from a local source; this commercial oil source is routinely used by present-day Ayurvedic practitioners. It would be important to consider variations in the properties of the oil based on the source of the raw seeds and processing conditions (63).

### Preparation of variations of the formulations

The recipes also did not call for any specific type of metal container for remedy preparation, and we used a stainless steel pot. Leaching of trace amounts of metals (silver, aluminium, copper) from containers could contribute to the efficacy of formulations (64). We also ‘modernized’ the preparation practices to use a mechanised grinder (as opposed to physical grinding with rock or stone), to ensure we obtained a uniform, consistent paste. This could influence the ‘release’ of ingredients into the plant paste, and use of a mortar and pestle could be an intermediate option. Similarly, the liquid component of the remedy recommends the use of water or milk, and we decided to use water. One way to account for these factors would be to prepare and test multiple variations of the traditional formulations.

### Sterilization of the formulations

Notably, the recipe did not call for any specific sterilization practices. This could be because the prolonged boiling process may itself have a sterilizing effect, or that microbes, if any, would be part of the ‘remedy’. It is important to consider the effects that modern-day sterilization practices (high temperature, filtration) might have on the constituents of the formulation. Accordingly, we did not sterilize the remedies, however, when incubated as controls, uninoculated sesame oil and traditional remedies did not demonstrate microbial growth (data not shown).

### Long term storage

We prepared and tested all the three formulations in a single batch. Sesame oil has high oxidative stability (65), and can be stored at room temperature for long periods of time. However, it would be important to consider possible deterioration in antimicrobial activity of the formulation, depending on the ingredients and their properties, and testing could be done in intervals of time.

### Testing the efficacy of traditional remedies with scientific assays and analyses

To test the anti-biofilm potential of reconstituted remedies we used standard microtiter-plate based biofilm assays. These are *in vitro* laboratory-based assays, in which thin biofilms form on the bottom and sides of polystyrene wells (39). Inspite of these obvious limiting factors, we chose to test the remedies with these assays for several reasons. Microtiter based biofilm assays can be used to rapidly test a large number of compounds and enable semi-quantitative estimations of biomass and metabolic activity of biofilms (39). In addition, a recent set of technical guidelines have been published to enable consistent testing and reporting (40). Therefore, these widely-employed assays lend well as ‘screening’ platforms. While these assays have been used to test various natural antimicrobial compounds, they are usually single ingredients and chemical extracts; here we show that these assays can be used to test ‘whole’ traditional remedies. It is important to consider that the medicinal preparations are in a form suitable for testing with these assays, for example, liquid preparations as opposed to a gel or cream. Moreover, the compatibility of the nature of the formulation, testing platform, and readout of the assays, is an important determinant. With the microtiter-based biofilm assays, the reconstituted remedies displayed differential activity, based on biomass formation and metabolic activity, against biofilm states of *P. aeruginosa*, a Gram negative pathogen, and *S. aureus*, which is Gram positive. This highlights the value of this assay as an initial screening tool for traditional remedies. Notably, these effects were seen against pre-formed, day-old biofilms as well as on the formation of new biofilms. This underscores the importance of testing a wide range of pathogens, and different biofilm states.

### Further evaluation and clinically-relevant testing

The biofilm inhibitory effects of the medicinal formulations could be attributed to the properties of ingredients, that include specific antimicrobial effects or interactions with the biofilm matrix (54,55,59,60). On the other hand, the possibility of a physical effect of the oil-based remedies, that prevents bacterial attachment or reduces the availability of oxygen is also possible (56). One study that employed crystal violet-based microtiter assays to evaluate the effects of essential oils (thyme, cinnamon, marjoram) on biofilm formation allowed the bacteria to attach to the substrate for 4 hours, after which the oils were added (56). The addition of certain oils resulted in a decrease in the number of attached cells, leaving mainly damaged cells (observed under SEM). Therefore, after an initial screening, the selected promising remedies can be further examined for the nature of their activity (bactericidal, bacteriostatic or matrix-modifying) as well as with *in vitro* systems that mimic infection sites and animal models (16,17). Though biological model systems are more clinically-relevant, they are not high-throughput, and require substantial expertise, making them unsuitable for high-throughput, preliminary testing.

### Historical and philosophical aspects

Other relevant considerations to developing traditional remedies include the need to establish open-minded teams, which include trained practitioners of ancient medical systems. Our team consisted of a doctor of modern medicine, a traditional medical practitioner, and scientists. This combined expertise made it possible to mine ancient treatises written in Sanskrit, translate them into English, and reconstitute, test and analyze them with modern scientific practice. It was certainly helpful that Ayurveda is still practiced and taught across India, and practitioners are familiar with these ancient treatises and preparations. It is also important to consider that traditional medicine is widely practiced in low-resource and underserved parts of the world (66,67), and as ‘ancientbiotics’ for biofilms gains traction as a field of study, it is important that researchers from these regions are included, and that tests and guidelines proposed are accessible in different settings.

Developing ‘ancientbiotics’ as anti-biofilm agents holds enormous potential, and there are a plethora of traditional remedies across historical systems of medicine to be explored (33,67–69). Is it evident there are several considerations and challenges involved in bringing these remedies out of treatise and to test their efficacy in the laboratory. While the exact details may vary across systems of medicine, historical texts, and preparation practices, this ‘treatise to test’ phase will include certain common themes. We believe that outlining them and discussing relevant considerations, will serve as a starting point for future studies along these lines.

## Author Contributions

V.M, S.K., A.B. and K.S.K. conceived and designed the study. A.B. and K.S.K. identified the traditional medicinal remedies. A.B. prepared the traditional medicinal remedies. V.M and S.K. did the experimental assays and statistical analyses. V.M, S.K. and K.S.K. analyzed the data and wrote the manuscript.

## Acknowledgements

We thank Sai Palande Datar, and Dr. Ambarish Khare, PhD, Department of Sanskrit, Tilak Maharashtra Vidyapeeth, Pune, for insightful discussions on the origins and practice of Ayurveda. We thank Dr. B. Balasubramanian, MBBS, MD, for assistance with Sanskrit translations.

