## Supplementary material for "Towards ‘Ancientbiotics’ for Biofilms: How can we bring traditional medicinal remedies out of treatise and into contemporary science?": Suppl Table 1

**Supplementary Files:**

**Table 1. Key words and their meanings from verses from historical medical treatises of Ayurveda**

| **Overall meaning of Ancient Text** | **Word in Sanskrit** | **Transliteration and Translation of key words in English** |
| --- | --- | --- |
| Figure 2A  *Sushruta Samhita*  Description of types of wounds  ‘There are two types of wounds. They are body wounds, because of internal disease states, and external wounds because of trauma, accidents, burns, and animal bites. Although both these classes of ulcers possess many features in common, they have been grouped under two  distinct heads on account of the diversity of their origin, the difference in remedial measures to be adopted in their treatment  and the variation in their strength and tenacity’. | व्रण | *Vrana* : Wound, Ulcer, Scratch |
|  | आगन्तु | *Agantu* : accidental |
|  | शारीर | *Sharira* : relating or belonging to or being in or produced from the body |
|  | पुरुष | *Purush :* Man |
|  | पशु | *Pashu :* Animal |
|  | पक्षि | *Pakshi :* Bird |
|  | व्याल | *Vyal :* Ferocious animal |
|  | सरीसृप | *Sarisrip :* Reptile |
|  | प्रहार | *Prahar :* Attack/blow/strike |
|  | अग्नि | *Agni :* Fire |

| Figure 2B  *Sushruta Samhita*  The problem of infected wounds  ‘The germination of worms due to flies in an ulcer is associated with extreme pain, swelling and bleeding, in case the worms eat  up the flesh. A decoction of certain herbs is an efficacious wash and healing medicine in such cases. The ulcer should be plastered with certain drugs, pasted with the urine of a cow, or washed with an alkaline wash (for expelling the vermin from it). As an alternative, the worms should be brought out of the ulcer by placing a small piece of raw flesh on the ulcer’. | मक्षिका | *Makshika* : Fly |
| --- | --- | --- |
|  | व्रणमागत्य | *Vranamagatya* : inside the wound |
|  | रुजा | *Ruja:* Pain |
|  | श्वयथु | *Shvayathu*: Swelling |
|  | रक्तस्राव | *Raktasrava*: Flow of blood |
|  | कृमिन् | *Krimin*: Having worms/affected with worms |

| Figure 2C  *Sushruta Samhita*  Principles of treatment of wounds and infected wounds  ‘The infected ulcer or sore should be washed with a decoction of select herbs cooked in oil and applied as a medicinal formulation on the wound for purification’. | दुष्टव्रण | *Dushtavrana* : non-healing wound |
| --- | --- | --- |
|  | शोधन | *Shodhana* : removal/cleansing |
|  | सुरस | *Suras* : holy basil |

| Figure 2D  *Bhavaprakash Nighantu*  Indian Materia Medica  Recommendation of the use of sesame oil for medicinal formulations  ‘Oils extracted from sesame increases stability of tissues, clears obstructed faeces and urine, is good for hair, skin and eyes, for treatment of fractures, as nasal drops, eye drops and ear treatments, and as medicated oil preparations’. | तिलतैल | *Teel Tail* : Sesame seed oil |
| --- | --- | --- |
|  | नस्य | *Nasya :* nose |
|  | कर्ण | *Karna* : ear |
|  | त्वच्य | *Tvachya :* conducive to health of skin |
|  | केश्य | *Keshya :* suitable for hair |
|  | व्रण | *Vrana :* wound |

| Figure 2E  *Dhravyagun Vignyan*  (Ayurvedic text book)  Use of *Bryophyllum pinnatum* or ‘*Parnabeej*’ for wound healing  ‘Parnabeej stops bleeding from the wound and has wound healing properties’. | पर्णबीज | *Parnabeej* : *Bryophyllum pinnatum* |
| --- | --- | --- |
|  | रक्त्तस्तम्भन | *Raktastanbhan :* stop bleeding |
|  | व्रणरोपण | *Vranaropana :* wound healing |

| Figure 2F  *Charaka Samhita*  *Cynodon dactylon* or ‘*Durva*’ for wound healing  ‘Extract or paste of Durva can be used to prepare oil. This oil has the property of wound healing’. | दूर्वा | *Durva* : *Cynodon dactylon* |
| --- | --- | --- |
|  | तैल | *Tail :* oil |
|  | व्रणरोपण | *Vranaropana :* wound healing |

| Figure 2G  *Bhavaprakash Nighantu*  Indian Materia Medica  Medicinal roles of ‘*Tulsi’* or ‘*Holy Basil*’  ‘Tulsi is also called Surasa. It stimulates digestion, and cures obstinate skin diseases such as leprosy, dysuria, pain in the sides of the chest. It also cures poisoning, parasitic infections, vomiting), asthma, piles and pain in the eyes.’ | तुलसी | *Tulsi* : *Ocimum tenuiflorum* |
| --- | --- | --- |
|  | सुरसा | *Surasa :* another name for Tulsi |
|  | कुष्ठ | *Kushta :* tenacious skin disease such as leprosy |

| Figure 2H  *Sharangdhar Samhita*  Preparation practices of herbal medicinal oils with 1:16:4 ratio of components and boiling process  ‘Count the paste of the herb. Get sesame oil four times the paste, and water or milk four times the oil’. | कल्क | *Kalka* : paste |
| --- | --- | --- |
|  | चतुर्गुण | *Chaturguna* : 4 times |
|  | द्रव्य | *Dravya* : Water |
|  | तैल | *Tail* : Oil |
